# Time-resolved dual root-microbe transcriptomics reveals early induced *Nicotiana benthamiana* genes and conserved infection-promoting *Phytophthora palmivora* effectors

**DOI:** 10.1101/098855

**Authors:** Edouard Evangelisti, Anna Gogleva, Thomas Hainaux, Mehdi Doumane, Frej Tulin, Clément Quan, Temur Yunusov, Kevin Floch, Sebastian Schornack

## Abstract

**Background:** Plant-pathogenic oomycetes are responsible for economically important losses on crops worldwide. *Phytophthora palmivora*, a broad-host-range tropical relative of the potato late blight pathogen, causes rotting diseases in many important tropical crops including papaya, cocoa, oil palm, black pepper, rubber, coconut, durian, mango, cassava and citrus.

Transcriptomics have helped to identify repertoires of host-translocated microbial effector proteins which counteract defenses and reprogram the host in support of infection. As such, these studies have helped understanding of how pathogens cause diseases. Despite the importance of *P. palmivora* diseases, genetic resources to allow for disease resistance breeding and identification of microbial effectors are scarce.

**Results:** We employed the model plant *N. benthamiana* to study the *P. palmivora* root infections at the cellular and molecular level. Time-resolved dual transcriptomics revealed different pathogen and host transcriptome dynamics. *De novo* assembly of *P. palmivora* transcriptome and semi-automated prediction and annotation of the secretome enabled robust identification of conserved infection-promoting effectors. We show that one of them, REX3, suppresses plant secretion processes. In a survey for early transcriptionally activated plant genes we identified a *N. benthamiana* gene specifically induced at infected root tips that encodes a peptide with danger-associated molecular features.

**Conclusions:** These results constitute a major advance in our understanding of *P. palmivora* diseases and establish extensive resources for *P. palmivora* pathogenomics, effector-aided resistance breeding and the generation of induced resistance to *Phytophthora* root infections. Furthermore, our approach to find infection relevant secreted genes is transferable to other pathogen-host interactions and not restricted to plants.

## Background

*Phytophthora* is a genus of plant-pathogenic oomycetes responsible for economically important losses on crops worldwide, as well as damage to natural ecosystems [1]. *Phytophthora infestans* is the causal agent of tomato and potato late blight in temperate climates and contributed to major crop losses during the Great Irish Famine [2]. *Phytophthora palmivora,* a broad-host-range tropical relative of *P. infestans* originating from south-eastern Asia [3] but now present worldwide due to international trade [4] causes root, bud and fruit rotting diseases in many important tropical crops such as papaya, cocoa, oil palm, black pepper, rubber, coconut, durian, mango, cassava and citrus [5-8]. In addition, *P. palmivora* infects roots and leaves of several model plant species such as *Medicago truncatula* [9], *Hordeum vulgare* [10] and *Arabidopsis thaliana* [11]. Despite its economic impact and widespread distribution, nothing is known about the molecular basis underlying its ability to infect many unrelated host species and the root responses associated with an infection.

*P. palmivora* has a hemibiotrophic lifestyle. Similar to other *Phytophthora* species, its asexual life cycle in plants is characterised by adhesion of mobile zoospores to the host tissue, encystment and germ tube formation [12]. Entry into the plant is achieved *via* surface appressoria and is followed by establishment of an apoplastic hyphal network. During this biotrophic stage *P. palmivora* projects haustoria into plant cells. These contribute to acquisition of nutrients and release virulence proteins known as effectors [13]. This is followed by a necrotrophic stage characterised by host tissue necrosis and the production of numerous sporangia which release zoospores [14].

Sequencing of *Phytophthora* genomes and transcriptomes has revealed repertoires of effector proteins that counteract plant defenses and reprogram the host in support of infection. Secretome predictions and subsequent evolutionary and functional studies have helped to understand how these pathogens cause diseases [15,16]. Oomycete effectors are secreted into the apoplast of infected plants. Some of them act inside plant cells and conserved RXLR or LFLAK amino acid motifs in their N-terminal parts have been associated with their translocation from the microbe into the host cell [17,18]. The LFLAK motif is present in Crinkler (CRN) effectors, named after a crinkling and necrosis phenotype caused by some CRN proteins when expressed in plants [19]. RXLR effectors are usually short proteins with little similarity to conserved functional domains in their C-termini. They localise to diverse subcellular compartments and associate with plant target proteins with key roles during infection [20].

Recent studies on bacterial and oomycete plant pathogens identified subsets of effectors that are conserved among a large number of strains. These so-called core effectors are responsible for a substantial contribution to virulence and thus cannot be mutated or lost by the pathogen without a significant decrease in virulence [21]. Thus, core effectors constitute highly valuable targets for identification of resistant germplasm and subsequent breeding disease-resistant crops [21–23].To date, the occurrence of such core effectors in oomycetes has largely been reported from plant pathogens with narrow economical host range such as *Hyaloperonospora arabidopsidis, Phytophthora sojae* [24] and *P. infestans* [25].

Plants have evolved a cell autonomous surveillance system to defend themselves against invading microbes [26]. Surface exposed pattern recognition receptors (PRRs) recognize conserved microbe-associated molecular patterns (MAMPs) released during infection, such as the *Phytophthora* transglutaminase peptide pep-13 [27,28]. In addition, plants are also able to recognize self-derived so-called damage-associated molecular patterns (DAMPs). These include intracellular peptides that get released in the apoplast upon wounding, such as systemins [29] and secreted plant peptides precursors with DAMP features that get processed in the apoplast [30–32]. Pathogen recognition initiates basal defense responses which include activation of structural and biochemical barriers, the MAMP-triggered immunity (MTI) [26]. Plant pathogens are able to overcome MTI by secreting effectors that suppress or compromise MTI responses, thereby facilitating effector-triggered susceptibility (ETS). In response, plants have evolved disease resistance proteins to detect pathogen effectors or effector-mediated modification of host processes, leading to effector-triggered immunity (ETI) [26]. *Phytophthora* genes encoding effectors which trigger a resistance response in host plants carrying the cognate disease resistance gene are often termed avirulence (AVR) genes. Cross-species transfer of PRRs and disease resistance genes against conserved MAMPs or AVR proteins has been successfully employed to engineer resistant crops [33,34].

Host cell responses to oomycete infections have mainly been studied in aboveground tissues and notably involve subcellular rearrangements of the infected cells, including remodelling of the cytoskeleton [14,35,36] and focal accumulation of secretory vesicles [37,38], which contribute to defense by delivering antimicrobial compounds to the extrahaustorial matrix [39,40]. Endocytic vesicles accumulate around oomycete haustoria [41] and the plant-specific small GTPase RAB5 is recruited at the extrahaustorial membrane during *Arabidopsis* infection by obligate biotrophs [42]. Several oomycete effectors target different stages of the host secretory pathway. In the apoplast, pathogen-secreted inhibitors have been associated with defense suppression. For instance, the apoplastic effector GIP1 from *P. sojae* inhibits the soybean endoglucanase EGaseA [43]. The *P. infestans* Kazal-like protease inhibitors EPI1 [44] and EPI10 [45] inhibit the *Solanum lycopersicum* defense protease P69B. The cystatin-like protease inhibitors EPIC1 and EPIC2B inhibit the cystein proteases PIP1 (*Phytophthora* Inhibited Protease 1) [46] and Rcr3 [47] as well as the papain-like protease C14 [48]. Interestingly, expression of the *P. infestans* RXLR effector AVRblb2 in plant cells prevents C14 protease secretion and causes an accumulation of protease-loaded secretory vesicles around haustoria [49].

In this study, we employ the model plant *N. benthamiana* [50] to study root infection by *P. palmivora.* Dual transcriptomics and *de novo* assembly of the *P. palmivora* transcriptome allowed us to define pathogen and plant genes expressed during the interaction. We identified major shifts in pathogen gene expression dynamics associated with lifestyle changes which, interestingly, are not mirrored by dramatic shifts in plant gene expression patterns. We characterised two conserved RXLR effectors, REX2 and REX3 that promote root infection upon expression in plants. We furthermore show that REX3 was able to interfere with host secretion. By studying host transcriptional changes upon infection we identified a gene encoding a secreted peptide precursor with potential DAMP motifs whose promoter was specifically activated at root tip infection sites. Hence, our work establishes a major resource for root-pathogen interactions, showcases examples of how to exploit these data, and provides inroads for effector-aided resistance breeding in tropical crops.

## Results

### *Phytophthora palmivora* exerts a hemibiotrophic lifestyle in *Nicotiana benthamiana* roots

To describe the infection development of the root pathogen *P. palmivora* we investigated the infection dynamics of hydroponically grown *N. benthamiana* plants root-inoculated with *P. palmivora* LILI-YKDEL [9] zoospores (**Figure 1**). Disease development was followed on the aerial parts (**Figure 1a**) since infected roots did not display visible disease symptoms. The plants looked healthy for up to 3 days (Symptom Extent Stage 1, SES 1). Disease progression in the aerial parts then resulted in a shrunken, brown hypocotyl and wilting of the oldest leaves (SES 2). This was rapidly followed by brown coloration and tissue shrinkage of the stem (SES 3) up to the apex (SES 4). Infected plants eventually died within 8 to 10 days (SES 5), indicating that *N. benthamiana* is susceptible to root infection by *P. palmivora* (**Figure 1a**).

**Figure 1.**
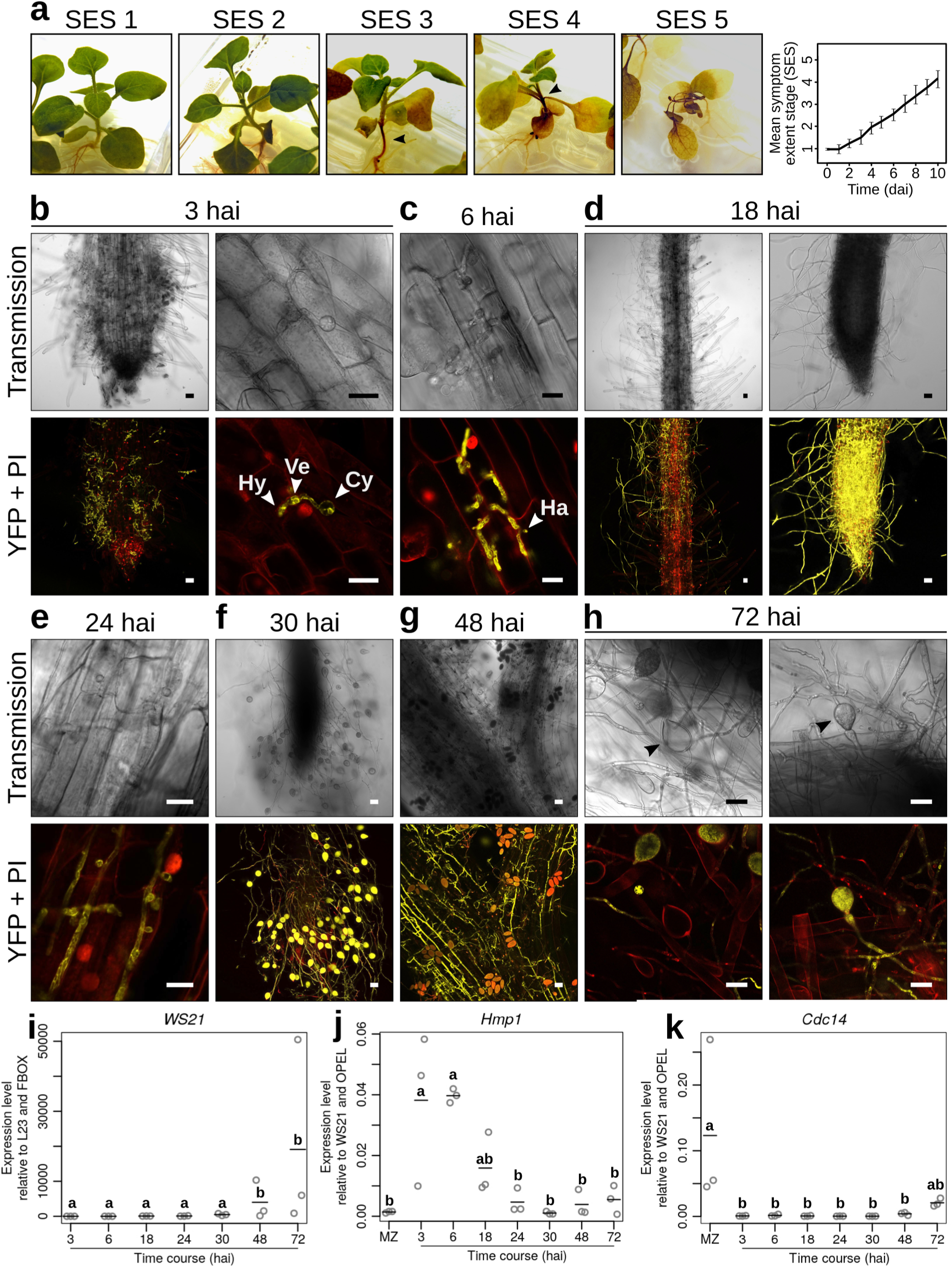
*Phytophthora palmivora* exerts a hemibiotrophic lifestyle in *Nicotiana benthamiana* roots. (**a**) Representative pictures of root-infected plantlets during *P. palmivora* infection, showing disease progression on the aboveground tissues. The successive symptom extent stages (SES) were used to define a disease index in order to quantitate disease progression over time. (**b-h**) Microscopic analysis of *N. benthamiana* roots inoculated with transgenic *P. palmivora* LILI expressing an endoplasmic reticulum (ER)-targeted YFP. Pictures were taken during penetration (**b**, 3 hours after inoculation (hai)), early infection (**c**, 6 hai), biotrophy (**d**, 18 hai and **e**, 24 hai), switch to necrotrophy (**f**, 30 hai) and necrotrophy (**g**, 48 hai and **h**, 72 hai respectively). Each pane shows transmission light (Transmission) and merged YFP fluorescence with propidium iodide (PI) staining (YFP + PI). Hy, hypha; Ve, vesicle; Cy: cyst; Ha: haustorium. Scale bar is 10 μm. (**i**) Quantification of *P. palmivora* biomass accumulation over time in *N. benthamiana* roots was measured by expression of *P. palmivora WS21* relative to *N. benthamiana L23* and *F-box* reference genes. (**j**, **k**) Expression of *P. palmivora* lifestyle marker genes *Hmp1* (k) and *Cdc14* (l) were measured over time relative to *P. palmivora WS21* and *OPEL* reference genes. Quantitative RT-PCR experiments were performed in triplicate. Dots represent values for each replicate. Bars represent the mean value. Statistical significance has been assessed using one-way ANOVA and Tukey’s HSD test (P < 0.05).

We next characterised the *P. palmivora* – *N. benthamiana* interaction on the microscopic level using the fluorescently labeled isolate LILI-YKDEL (**Figure 1b-h**). Infection events were observed at 3 hours after inoculation (hai). Zoospores were primarily attracted to root tips, where they encysted and germinated. Appressoria were differentiated at this stage and, when infection of the first cell had already occurred, an infection vesicle and subjacent nascent hyphae were also observed (**Figure 1b**). Haustoria were visible from 6 hai - 24 hai, indicative of biotrophic growth (**Figure 1c-e**). At 18 hai, *P. palmivora* hyphae grew parallel to the cell files in the root cortex, forming a clear colonisation front between infected and non-infected tissues. In addition, extraradical hyphal growth was observed near the root tip (**Figure 1d**). First sporangia occurred at 30 hai (**Figure 1f**). Consistent with the symptoms observed on aerial parts, hypocotyl colonization occurred between 30 hai and 48 hai (**Figure 1g**). Finally, the presence of empty or germinating sporangia at 72 hai suggests possible secondary infections (**Figure 1h**). Therefore, *P. palmivora* asexual life cycle completes within 72 hai in *N. benthamiana* roots.

We supported our microscopic studies with biomass quantification based on transcript levels of the *P. palmivora* 40S ribosomal protein S3A (WS21) (**Figure 1i**). We further characterized the different stages observed microscopically by quantifying expression of the *P. infestans* orthologs of *Hmp1* (haustorium-specific membrane protein) [51] (**Figure 1j**) and the cell-cycle regulator *Cdc14* [52] (**Figure 1k**). *Hmp1* transcripts peaked between 3 hai and 6 hai and then decreased at later stages. By contrast, *Cdc14* transcripts increased at late time points (48 hai and 72 hai). Taken together, these results further support the conclusion that *P. palmivora* exerts a hemibiotrophic lifestyle in *N. benthamiana* roots.

### *De novo* assembly of *P. palmivora* transcriptome from mixed samples

We performed dual sequencing permitting *de novo* assembly of a *P. palmivora* transcriptome as well as an assessment of transcriptional changes in both, host and pathogen over time. We extracted RNA from infected and uninfected *N. benthamiana* roots at six time points matching the key steps of the interaction identified by microscopy: 6 hai, 18 hai, 24 hai, 30 hai, 48 hai and 72 hai and an axenically grown *P. palmivora* sample containing mycelia and zoospores (MZ). Using Illumina HiSeq 2500 paired-end sequencing we obtained a relatively uniform read depth of 50-60 M reads per sample (**Table S1**). To cover all possible transcripts we reconstructed the *P. palmivora* transcriptome *de novo,* combining *ex planta* and *in planta* root samples as well as 76 nt Illumina paired-end reads from infected *N. benthamiana* leaf samples (more than 515 M reads, **Figure 2a**, **Table S1**).

**Figure 2.**
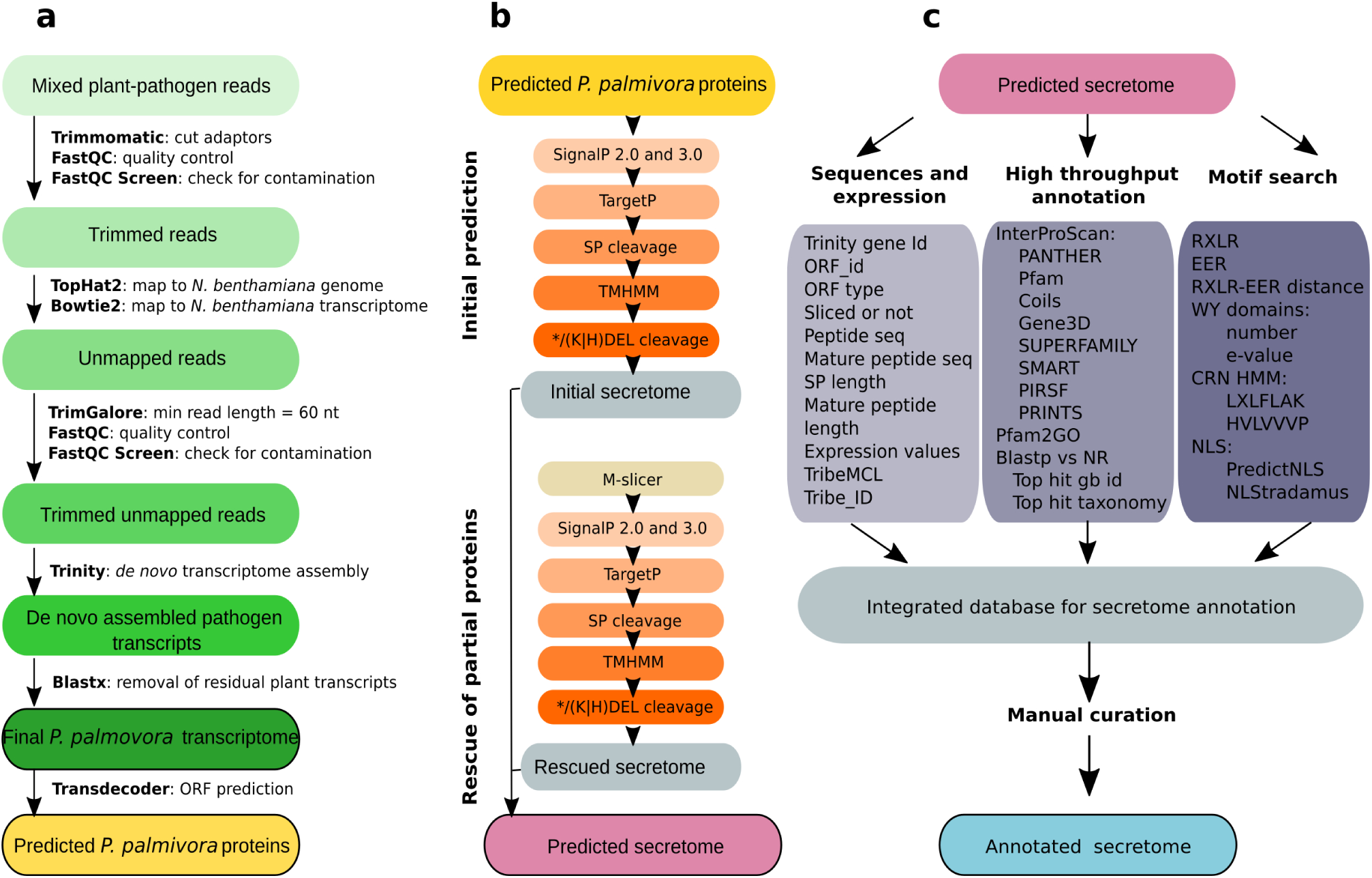
Overview of *P. palmivora* sequencing data analysis workflows. (**a**) Selection of *P. palmivora* reads from mixed samples and *de novo* assembly of transcriptome. (**b**) Secretome prediction. (**c**) Pipeline for automated secretome annotation. Final product of each pipeline are highlighted by bold lines. Abbreviations: SP: signal peptide; NLS: nuclear localization signal; CRN: crinkler.

Following standard adaptor trimming and read quality control, we applied a two-step filtering procedure (**Figure 2a**) to separate pathogen reads from plant host reads. First we mapped the pooled read dataset to the *N. benthamiana* reference genome and collected unmapped read pairs. Recovered reads were subsequently mapped to the *N. benthamiana* transcriptome [53]. Reads not mapped to either host plant genome or transcriptome were used to run assemblies. Short reads (<60 nt) were filtered out to produce transcripts of better quality and coherence. Final *de novo* Trinity assemblies were run from 190 M pre-processed, properly paired and cleaned reads ( **Table S1**). This yielded 57’579 ‘trinity genes’ corresponding to 100‘303 transcripts with an average backwards alignment rate of 76%, indicative of an overall acceptable representation of reads and therefore reasonably good assembly quality [54]. 9’491 trinity genes (20’045 transcripts including all isoforms) were removed by additional checks for residual plant contamination, resulting in a final *P. palmivora* transcriptome of 48’089 trinity genes corresponding to 80’258 transcripts (**Table 1).** We further selected 13'997 trinity genes (corresponding to 27106 transcripts) having the best expression support (**Supplementary Dataset 1**).

We assessed completeness of *P. palmivora* assembly by benchmarking nearly universal single-copy orthologs (BUSCO) [55] (**Table 1**) and compared to the BUSCO content of *P. infestans*, *P. sojae* and *P. parasitica* transcriptomes. We identified 326 BUSCO genes (76% of eukaryotic BUSCO genes) in our *P. palmivora* assembly, 348 (81%) in *P. infestans,* 343 (80%) in *P. sojae* and 360 (84%) in *P. parasitica.* (**Table 1**, **Supplementary Figure 1**). We also surveyed 14 publicly available *Phytophthora* genomes, yielding 20 additional BUSCO genes absent from all transcriptomes. Interestingly, the remaining 35 BUSCO genes were consistently missing from all analysed *Phytophthora* genomes and transcriptomes (**Supplementary Table 2**). These results suggest that our *P. palmivora* (LILI) transcriptome assembly actually contained 87% of BUSCO genes occurring in *Phytophthora* Hence, our assembly shows acceptable quality and integrity and can be used as a reference for further studies.

**Table 1.** *De novo* transcriptome assembly statistics for *P. palmivora*

| **Metric** | **Value** |
| --- | --- |
| **General assembly stats** |  |
| Total trinity 'genes' | 48'088 |
| Total trinity transcripts | 80'258 |
| Percent GC | 48.27 |
| Smallest contig | 201 |
| Largest contig | 217'111 |
| **Stats based on all transcript contigs:** |  |
| Contig N10 | 6'335 |
| Contig N20 | 4'510 |
| Contig N30 | 3'443 |
| Contig N40 | 2'621 |
| Contig N50 | 1'978 |
| Median contig length | 282 |
| Average contig length | 788 |
| Total assembled bases | 63'247'196 |
| **Stats based on the longest isoform per gene:** |  |
| Contig N10 | 6'738 |
| Contig N20 | 4'702 |
| Contig N30 | 3'604 |
| Contig N40 | 2'724 |
| Contig N50 | 2'003 |
| Median contig length | 273 |
| Average contig length | 765 |
| Total assembled bases | 36'825'977 |
| **Eukaryotic BUSCO genes** |  |
| Complete single-copy BUSCOs | 256 |
| Complete duplicated BUSCOs | 40 |
| Fragmented BUSCOs | 30 |
| Missing BUSCOs | 103 |
| Total BUSCO groups searched | 429 |
| Total BUSCO genes recovered | 326 (76%) |
| *Phytophthora* BUSCOs | 374 |
| Updated % recoverable BUSCOs | 87 |

### Clustering of plant and pathogen samples reflects different temporal dynamics during infection

To explore temporal expression dynamics of plant and pathogen genes we separately mapped initial reads back to the reference *N. benthamiana* transcriptome (https://solgenomics.net/) as well as to our *P. palmivora* transcriptome assembly (**Supplementary dataset 2**, **Supplementary dataset 3**). Principal component analysis (PCA) of plant samples revealed a major difference between infected and uninfected samples (91% of variance; **Figure 3a**). Plant transcript profiles from infected samples could be further assigned into three groups: 6 hai; 18-24-30 hai; and 48-72 hai (4% of variance; **Figure 3a**). Conversely, PCA analysis of *P. palmivora* transcript profiles identified two groups corresponding to early infection (6 to 24 hai) and late infection with MZ (48 and 72 hai), while 30 hai was kept apart (66% of variance; **Figure 3b**). Taken together, these results suggest different behaviour of plant and pathogen transcript profiles at the same times post infection.

**Figure 3.**
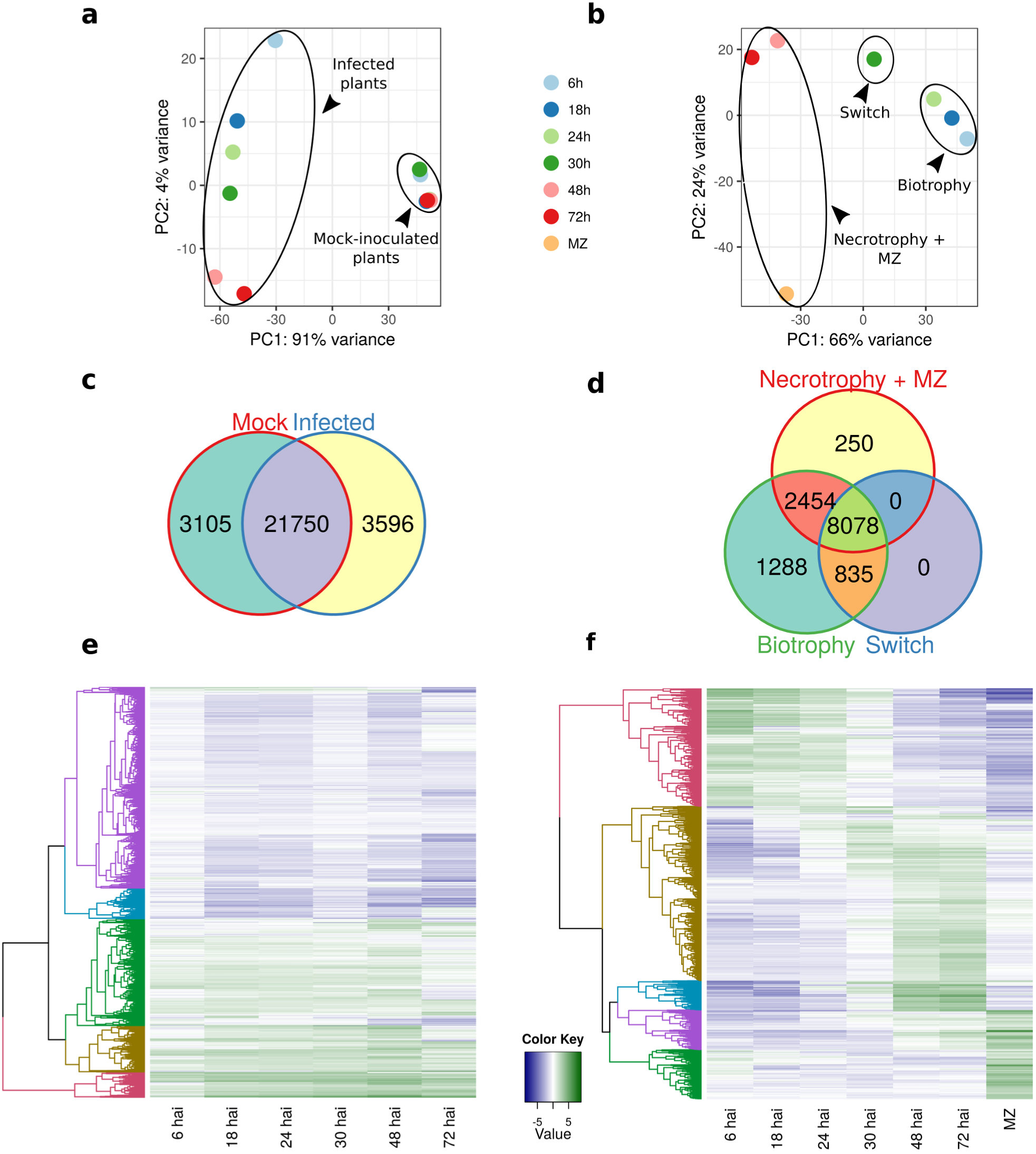
*N. benthamiana* and *P. palmivora* transcriptomes show different temporal dynamics during interaction. (**a-b**) PCA clustering of full transcriptional profiles of *N. benthamiana* (**a**) and *P. palmivora* (**b**). (**c-d**) Venn diagrams show shared genes expressed in groups identified by PCA analysis for *N. benthamiana* (**c**) and *P. palmivora* (**d**). Genes with TPM ≥ 5 were considered to be expressed. (**e-f**) Hierarchical clustering of major classes of differentially expressed genes (p-value < 10-3, LFC ≥ 2) in *N. benthamiana* (**e**) and *P. palmivora* (**f**) transcriptomes. Relative expression levels of each transcript (rows) in each sample (column) are shown. TPMs were log2-transformed and then median-centered by transcript. Plant samples were centered according to the full set of mock and infected samples, only infected samples are shown. Abbreviation: MZ: axenically grown mycelium with sporangia.

We identified 6590 plant and 2441 pathogen differentially expressed genes (DEGs) by performing differential expression analysis (LFC ≥ 2, FDR *p* <10-3) on all possible sample pairs (**Figure 3e**, **f**, **Figure S3**). Hierarchical clustering revealed 236 *P. palmivora* genes upregulated exclusively during biotrophy (from 6 to 30 hai), while all other stages shared sets of induced and expressed genes (**Figure 3f**, **d**). Interestingly, major shifts in expression patterns occurred at 30 hai. Taken together with PCA grouping, this result suggest that 30 hai represents a transition stage from a biotrophic to a necrotrophic lifestyle.

In contrast to the pathogen, the plant transcriptome did not undergo sharp transitions over time and was instead characterised by steady up- or downregulation (**Figure 3e**, **c**). Therefore, we utilised repeated upregulation of a gene in at least two timepoints as selective criterion to alleviate the absence of replicates resulting in 2’078 up and 2’054 downregulated genes. From these we validated 5 out of 6 genes with low or no expression under control conditions and high expression levels during infection using qRT-PCR (**Figure S10**). GO term analysis revealed that upregulated genes are enriched in biological processes related to hormone metabolism, abiotic stress (including oxidative stress, response to heat and wounding), defense, biosynthesis, transport, regulation of transcription and protein modification by phosphorylation and ubiquitination (**Supplementary Dataset 4**). Notably, we detected upregulation of numerous ethylene-responsive transcription factors (ERFs), indicating reprogramming of stress-specific defense regulation. Representatives of significantly enriched GO categories relevant for defense response include genes encoding endopeptidase inhibitors, such as Kunitz-type trypsin inhibitors. We also found upregulation of 48 genes encoding O-glycosyl hydrolases. In addition, we detected upregulation of trehalose biosynthesis pathway genes. Conversely, down-regulated genes showed overall enrichment in biological processes associated with photosynthesis, cellulose biosynthesis and cell division. Taken together, these results suggest that infected *N. benthamiana* roots undergo major transcriptional and post-translational reprogramming leading to an overall activation of stress and defense responses.

### *P. palmivora* secretome prediction and annotation identifies a set of effector candidate genes

Pathogen-secreted effectors and hydrolytic enzymes are hallmarks of *Phytophthora* infection [56]. Therefore, we probed our *P. palmivora* transcriptome for transcripts encoding secreted proteins. A TransDecoder-based search for candidate open reading frames (ORFs) [57] identified 123'528 ORFs from predicted trinity genes (isoforms included). We then analyzed the predicted ORFs using an automated pipeline for secretome prediction (**Figure 2b**) building on existing tools [58–60]. The pipeline was designed to predict signal peptides and cellular localisation with thresholds specific for oomycete sequences [61,62] and to exclude proteins with internal transmembrane domains and/or an endoplasmic reticulum (ER) retention signal. We identified 4’163 ORFs encoding putative secreted proteins.

Partial translated ORFs which were not predicted as secreted were subjected to an additional analysis (M-slicer) (**Figure 2b**) and resubmitted to the secretome prediction pipeline. This improved procedure allowed us to rescue 611 additional ORFs encoding putative secreted proteins. In total, we identified 4’774 ORFs encoding putative secreted *P. palmivora* proteins. We further selected a single representative secreted ORF for genes with sufficient expression support (TPM ≥ 1 in 3 or more samples). This yielded 2’028 *P. palmivora* genes encoding putative secreted proteins (**Supplementary Dataset 5**).

To maximise functional annotation of the *P. palmivora* secretome we used an integrative approach (**Figure 2c**) tailored to the use of known short motifs characteristic of oomycete secreted proteins. The pipeline contains three major blocks. The first block integrated all the sequence information, assignment to 2’028 non-redundant genes encoding secreted proteins as well as expression data. The second block combines results of homology searches, for both full-length alignments (blastn and blastx) and individual functional domains (InterProScan). The third block was designed to survey for known oomycete motifs and domains (such as RXLR, EER, WY for RXLR-effectors; LXLFLAK for crinklers and NLS for effectors in general). The pipeline produced an initial secretome annotation (**Figure 2c**) which was then manually curated to avoid conflicting annotations. This strategy allowed us to to assign a functional category to 768 (38 %) of predicted secreted proteins (**Table 2**).

Amongst predicted cytoplasmic effectors the most prominent category encompasses 140 RXLR effectors. Of these 123 have a conserved RXLR motif followed by dEER motif. WY-domains were found in 30 RXLR-EER effectors and 3 RXLR effectors. Some RXLR effectors are unusually long (> 400 a.a; average length of RXLR effectors being 204 aa) suggesting multiple effector domains linked together. For instance, the effector domain of PLTG_07082 consisted of 8 internal repeats of a WY-domain. It remains to be tested whether multiple WY domains within one effector fulfil different and independent roles.

PFAM searches revealed one full length RXLR effector protein (PLTG_09049) carrying a C-terminal NUDIX domain. PFAM predictions assigned to partial genes identified two putative effectors bearing a NUDIX domain PF00293 (PLTG_05400) and a MYB/SANT domain PF00249 (PLTG_06121).

Sequence similarity searches for RXLR effectors matching known oomycete avirulence proteins revealed PLTG_13552 as being similar to *P. infestans* AVR3a (PiAVR3a) (**Supplementary Figure 2**). Notably, *P. palmivora* AVR3a (PLTG_13552) harbours the K80/I103 configuration, but combined with a terminal valine instead of a tyrosine in PiAVR3a [63]. It thus remains to be tested whether PLTG_13552 is capable of triggering a *R3a*-mediated hypersensitive response.

Our pipeline only identified 3 genes encoding putative CRN effectors (PLTG_06681, PLTG_02753, PLTG_03744). Crinklers often lack predictable signal peptides, but instead might be translocated into plant cells by an alternative mechanism [64]. An independent survey using HMM-prediction without prior signal peptide prediction revealed a total of 15 CRN motif-containing proteins. Notably, the putative CRN effector PLTG_06681 carries a C-terminal serine/threonine kinase domain (PF00069) and shows low sequence similarity (34%) to *P. infestans* effector CRN8 [65].

The *P. palmivora* secretome also contained a substantial number of apoplastic effectors (**Table 2**). We identified 28 genes encoding extracellular protease inhibitors, including extracellular serine protease inhibitors (EPI) with up to five recognisable Kazal domains, several cystatins and cysteine protease inhibitors (EPICs) (**Supplementary Dataset 5**). PLTG_05646 encodes a cathepsin protease inhibitor domain followed by a cysteine protease and an ML domain (PF02221, MD-2-related lipid recognition domain). We also identified 28 proteins with small cysteine rich (SCR) signatures, 18 of them being encoded in full-length ORFs, but only six where the mature peptide is shorter than 100 aa. Longer SCRs can harbour tandem arrangements (PLTG_08623). In one case an SCR is linked to a N-terminal PAN/APPLE domain, which is common for carbohydrate-binding proteins [66].

Additionally the *P. palmivora* secretome contains 90 proteins carrying potential MAMPs, including necrosis-inducing proteins (NLPs), elicitins and lectins. Out of 24 NLPs, four (PLTG_05347,PLTG_07271, PLTG_13864, PLTG_01764) carry a pattern of 20 amino acid residues which is similar to the immunogenic nlp20 motif (AiMYySwyFPKDSPVTGLGHR, less conserved amino acids in lower case) [67]. Transcripts encoding elicitins and elicitors in the *P. palmivora* secretome belong to the group of highest expressed ones during infection (**Supplementary Dataset 5**). We identified six transglutaminases, five of them (PLTG_04342, PLTG_02581, PLTG_10032, PLTG_10034 and PLTG_10033) carrying a conserved pep13-motif [28].

Taken together, *de novo* transcriptome assembly followed by multistep prediction of ORF encoding potentially secreted proteins and a semi-automated annotation procedure allowed us to identify all major classes of effectors characteristic to oomycetes as well as *P. palmivora*-specific effectors with previously unreported domain arrangements. Our data suggest that *P. palmivora*’s infection strategy relies on a diverse set of extracellular proteins many of which do not match to previously characterised effectors.

**Table 2.**
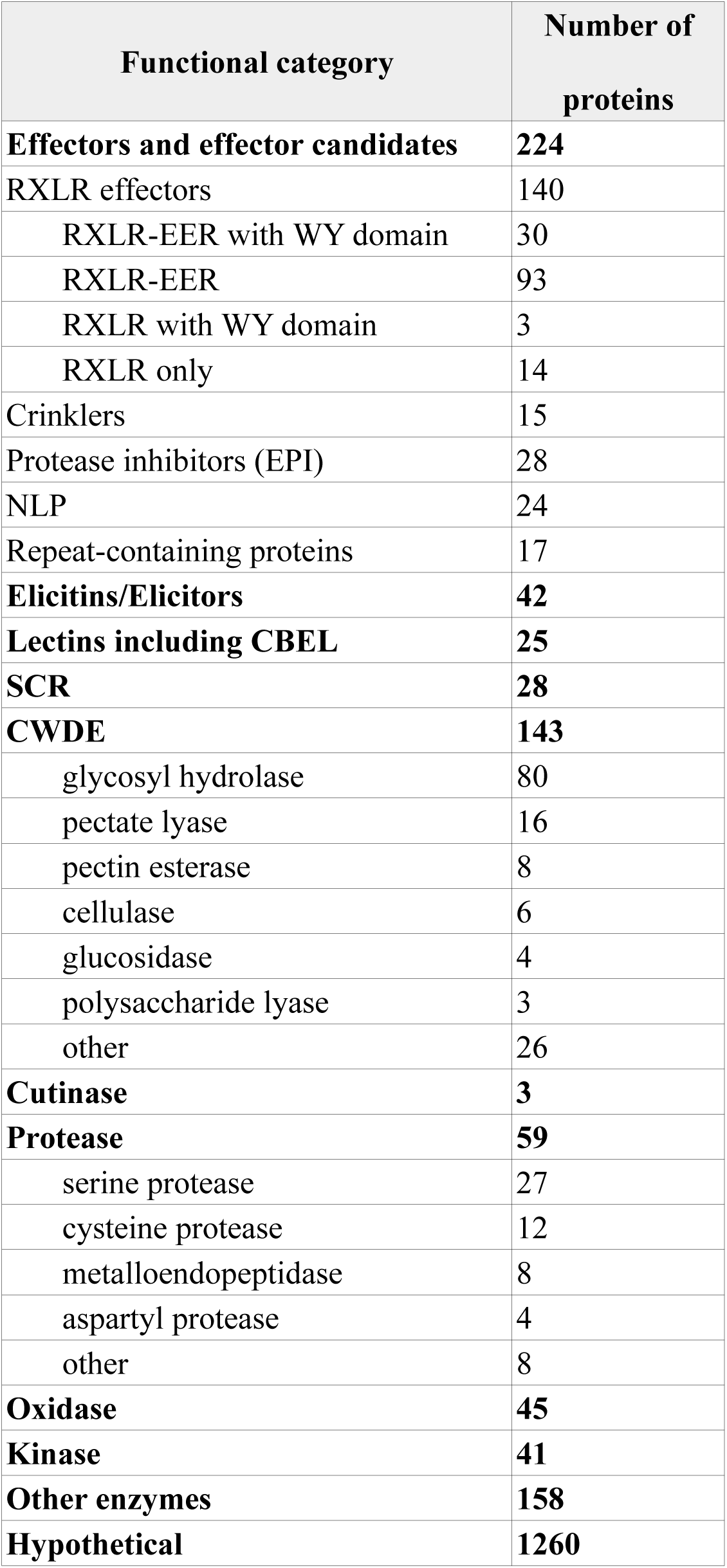
Representation of classes putative extracellular proteins in *P. palmivora* secretome (strain LILI)

| Functional category | Number of proteins |
| --- | --- |
| **Effectors and effector candidates** | **224** |
| RXLR effectors | 140 |
| RXLR-EER with WY domain | 30 |
| RXLR-EER | 93 |
| RXLR with WY domain | 3 |
| RXLR only | 14 |
| Crinklers | 15 |
| Protease inhibitors (EPI) | 28 |
| NLP | 24 |
| Repeat-containing proteins | 17 |
| **Elicitins/Elicitors** | **42** |
| **Lectins including CBEL** | **25** |
| **SCR** | **28** |
| **CWDE** | **143** |
| glycosyl hydrolase | 80 |
| pectate lyase | 16 |
| pectin esterase | 8 |
| cellulase | 6 |
| glucosidase | 4 |
| polysaccharide lyase | 3 |
| other | 26 |
| **Cutinase** | **3** |
| **Protease** | **59** |
| serine protease | 27 |
| cysteine protease | 12 |
| metalloendopeptidase | 8 |
| aspartyl protease | 4 |
| other | 8 |
| **Oxidase** | **45** |
| **Kinase** | **41** |
| **Other enzymes** | **158** |
| **Hypothetical** | **1260** |

### Most differentially expressed secreted proteins have their highest expression during biotrophy

In order to highlight dynamic expression changes of *P. palmivora* genes during infection, we performed fuzzy clustering of *P. palmivora* DEGs (**Figure 4**) to lower sensitivity to noisy expression signals and to distinguish between expression profiles, even if they partially overlapped [68]. We identified 12 expression clusters falling into four main groups according to their temporal expression level maximum (**Figure 4a**). Group A was composed of 2 clusters containing genes down-regulated during infection. By contrast, expression levels of genes from group B peaked during biotrophy (6-24 hai). Group C was composed of 2 clusters of genes for which transcripts accumulated mostly at 30 hai, while group D was formed of four clusters of genes with maximum expression during necrotrophy (48, 72 hai). Group B showed an overall enrichment in all classes of genes encoding secreted proteins (**Figure 4b**) while groups A and C were enriched in elicitin-encoding genes. SCRs were enriched in group D. Also in group D and characterised by strong transcriptional induction was a gene (PLTG_02529) encoding several repetitions of an unknown *Phytophthora*-specific amino acid motif. Expression dynamics of 18 *P. palmivora* genes from different clusters were validated by qRT-PCR. Fourteen genes displayed expression patterns consistent with the results of *in silico* prediction (**Figure S4b-o**). Taken together, these results suggest that *P. palmivora* transcriptome dynamics reflect the main lifestyle transitions observed by microscopic analysis of the infection process, and that a major upregulation of secreted proteins occurs during biotrophy in agreement with the occurrence of haustoria, which are a major site for pathogen secretion [13].

**Figure 4.**
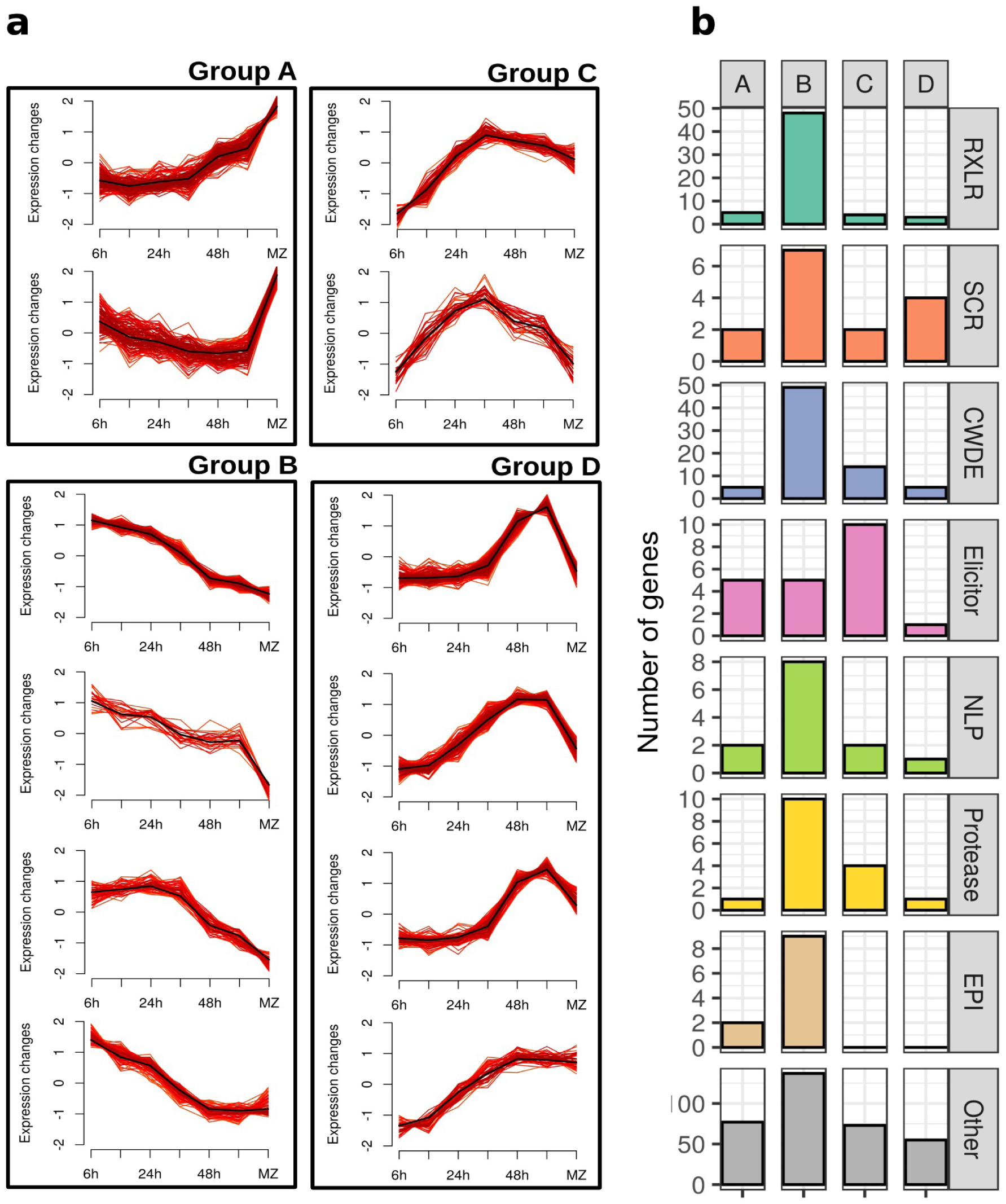
Temporal dynamics of *P. palmivora* differentially expressed genes (DEGs) during infection time course. Fuzzy clustering was performed on *P. palmivora* DEGs. Only genes with cluster membership values ≥ 0.7 are shown, *i.e.* alpha cores (**a**). Functional distribution of secreted proteins for the grouped clusters is shown in (**b**). Abbreviations: RXLR: RXLR-effector; SCR: small cysteine-rich protein; CWDE: cell wall degrading enzyme; NLP: necrosis inducing protein; EPI: protease inhibitor; Other: other genes encoding proteins predicted to be secreted without specific functional category assigned.

### Conserved RXLR effectors among *P. palmivora* isolates confer enhanced plant susceptibility to root infection

We next focussed on the characterisation of four RXLR effectors upregulated during infection (**Figure S4**) and named them *REX1* (PLTG_01927; GenBank accession KX130348), *REX2* (PLTG_00715; GenBank accession KX130350), *REX3* (PLTG_00687; GenBank accession KX130351) and *REX4* (PLTG_13723; GenBank accession KX130352). REX1-4 sequences from *P. palmivora* isolates with diverse geographic and host species origin (**Table S4**) were obtained by PCR and amplicon sequencing. Primers specific for *REX1-4* generated amplicons from at least 13 of the 18 isolates (*REX1*: 15, *REX2*: 15, *REX3*: 16, *REX4*: 13, **Figure S5**) encoding proteins with high levels of amino acid sequence conservation. In particular *REX2* and *REX3* were almost invariant, with one and two amino acid substitutions, respectively (**Figure S6**).

N-terminal translational GFP fusions of FLAG-tagged *REX* coding sequences (referred to as GFP:FLAG-REX1-4) expressed in roots of stable transgenic *N. benthamiana* plants (**Figure 5**, **Figure S7**) or transiently in the leaf epidermis (**Figure S8a-d**) showed nuclear and cytoplasmic fluorescence at 24 hai originating from expression of full length GFP:FLAG-REX1,2 and REX4 protein fusions (**Figure S8e**). In contrast to the other three, GFP:FLAG-REX3 fluorescence signals were much weaker in the leaf epidermis nucleus compared the cytoplasmic signals and absent from root nuclei (**Figure 5c**, **Figure S8c**).

**Figure 5.**
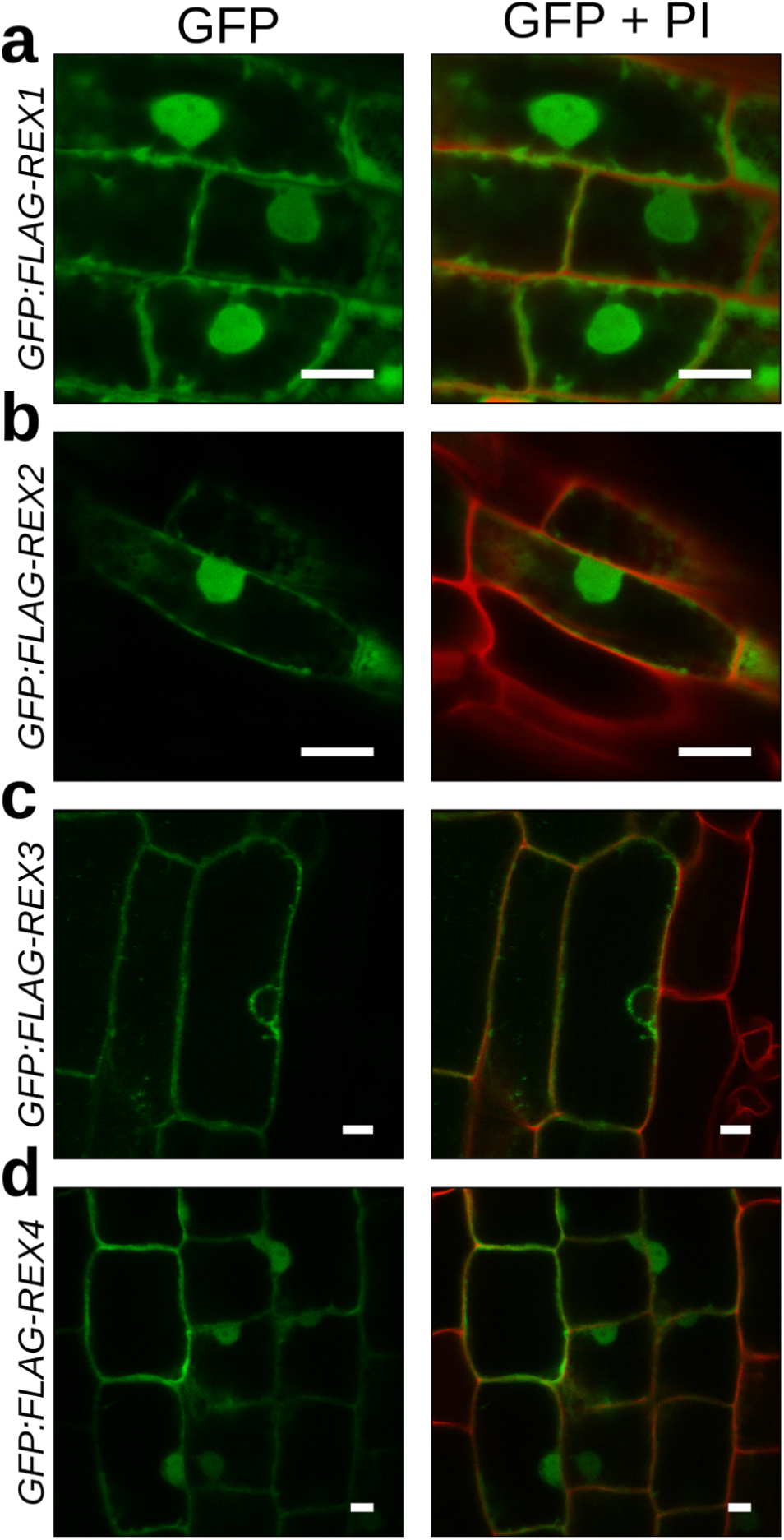
Spatial distribution of REX effectors in *N. benthamiana* roots. (**a-d**) Transgenic *N. benthamiana* plants expressing GFP:FLAG-REX fusion proteins were regenerated from leaf explants and grown to seeds. Subcellular localisation of GFP:FLAG-REX1-4 was assessed on seedling roots stained with propidium iodide (PI). GFP:FLAG-REX1 (**a**), GFP:FLAG-REX2 (**b**) and GFP:FLAG-REX4 (**d**) accumulated in the cytoplasm and in the nucleus. GFP:FLAG-REX3 (**c**) was detected in the cytoplasm but was excluded from the nucleus. Scale bar is 10 μm.

To determine the contribution of REX1-4 to *N. benthamiana* root infection, we then challenged hydroponically grown transgenic plants expressing GFP:FLAG-REX1-4 or GFP16c-expressing plants (ER-targeted GFP) with *P. palmivora* zoospores (**Figure 6a**,**b**) and monitored disease progression into aerial tissues over time using a disease index ranking from 1 to 5 derived from the symptoms previously reported (**Figure 1**). Transgenic plants expressing GFP:FLAG fusions of the highly conserved REX2 and REX3 effectors displayed significantly accelerated disease symptom development (P-values of 5.4 10–16 and 0.013, respectively) compared to GFP16c control plants, while expression of GFP:FLAG-REX1 and GFP:FLAG-REX4 did not enhance susceptibility (*P*-values of 0.66 and 0.24, respectively) (**Figure 6a**,**b**).

**Figure 6.**
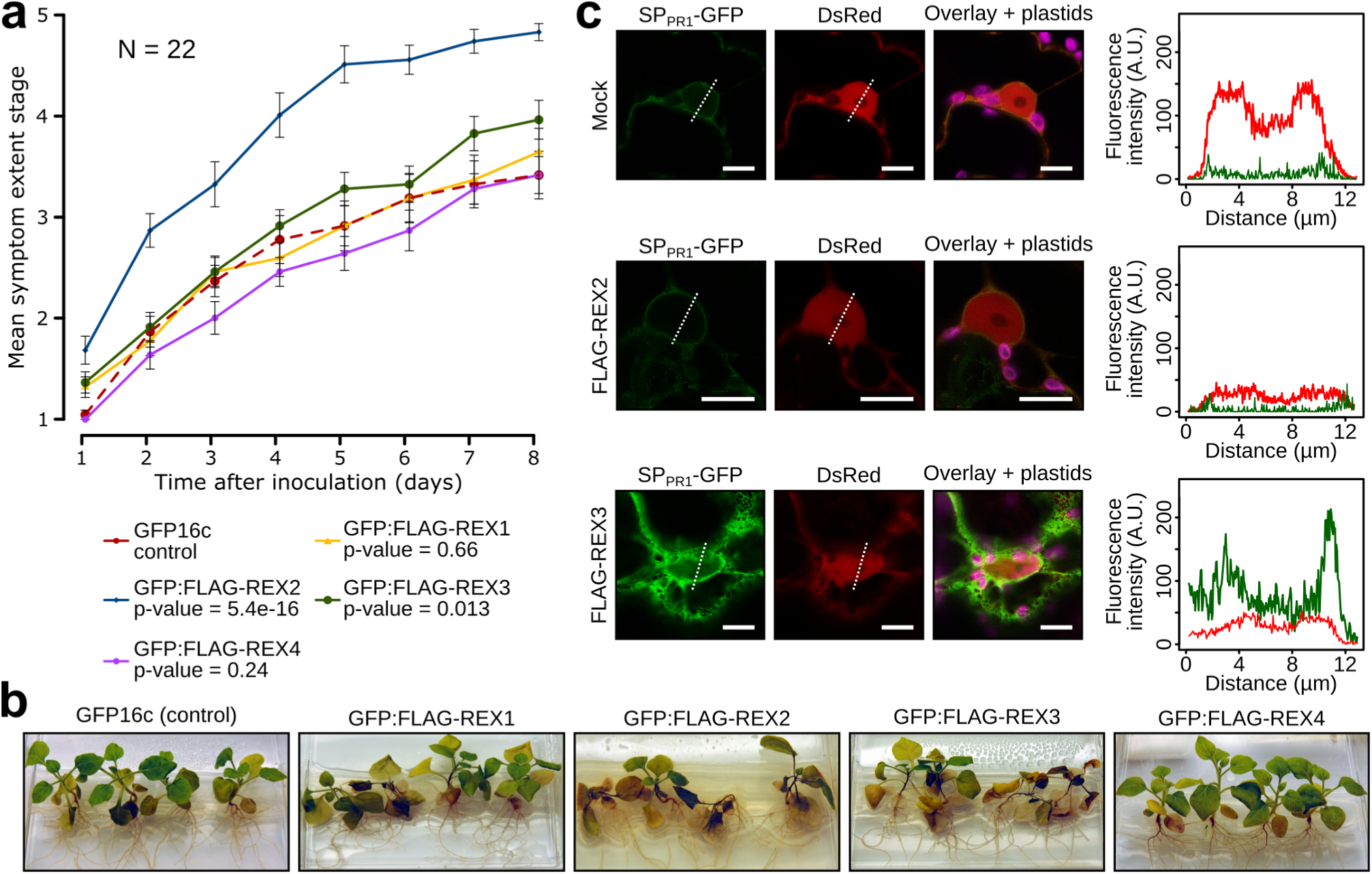
REX2 and REX3 increase *N. benthamiana* susceptibility to *P. palmivora* and REX3 interferes with host secretion. Transgenic *N. benthamiana* plants expressing GFP16c (control) or GFP:FLAG-REX1 to GFP:FLAG-REX4 were challenged with zoospores from *P. palmivora* YKDEL and disease progression was ranked over time using the previously defined symptom extent stages (SES). (**a**) Representative disease progression curves for transgenic plants expressing GFP:FLAG-REX1 (yellow), GFP:FLAG-REX2 (blue), GFP:FLAG-REX3 (green) or GFP:FLAG-REX4 (magenta), when compared to GFP16c control plants (red dashed). P-values were determined based on Scheirer–Ray–Hare nonparametric two-way analysis of variance (ANOVA) for ranked data. The experiment was carried out in duplicate (N = 22 plants). (**b**) Representative pictures of infected plants, 8 days after infection. (**c**) Disease-promoting effectors REX2 and REX3 were coexpressed with a secreted GFP construct (SPpr1-GFP) in *N. benthamiana* leaves. GFP fluorescence was quantified along the nucleus.

**Figure 7.**
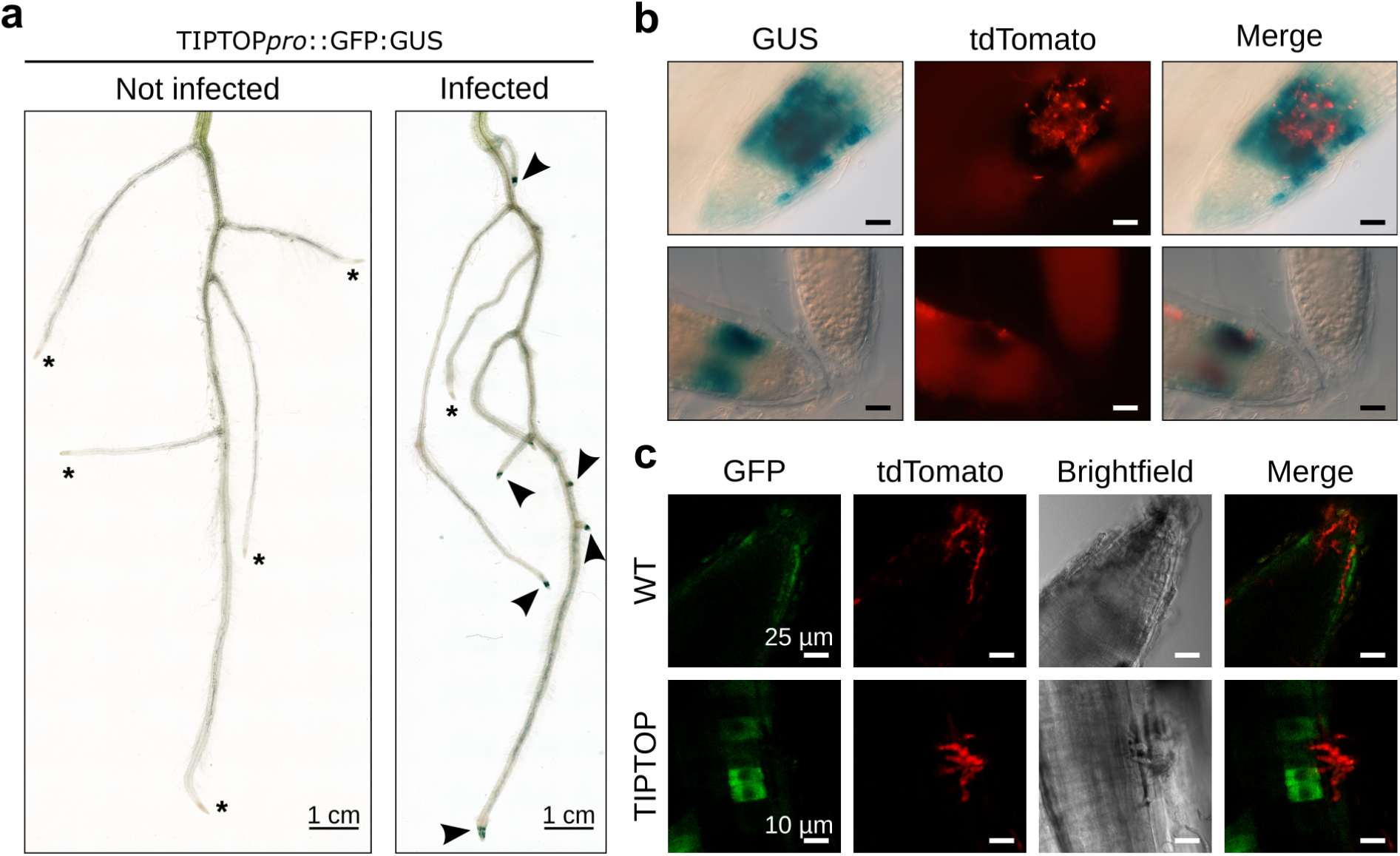
The promoter of a gene encoding the secreted peptide TIPTOP is upregulated during early biotrophy in *N. benthamiana* roots. (**a**) Representative pictures of GUS stained whole root systems of *N. benthamiana* transgenics carrying TIPTOP*pro*⸬GFP:GUS, non infected or 16 hours after infection with *P. palmivora* LILI-tdTomato. Stars represent unstained root tips. Arrowheads represent stained root tips. (**b**) Representative pictures of infected root tips after GUS staining, showing GUS signal at the vicinity of infection sites (top pane). Uninfected root tips from the same plant do not show any staining (bottom pane). Scale bar is 25 μm. (**c**) Representative pictures of GFP signal at the root tip of infected *N. benthamiana* transgenics expressing GFP:GUS fusion under the control of *TIPTOP* promoter.

### REX3 impairs plant secretion processes

Suppression of defense component secretion has previously been found to be targeted by at least two mechanisms [48,49]. We thus investigated the ability of the infection promoting REX2 and REX3 effectors to suppress host secretion (**Figure 6c**). We generated pTrafficLights, a vector which enables expression of a secreted GFP (SPPR1-GFP) together with a nuclear-cytoplasmic DsRed from the same *A. tumefaciens* T-DNA (**Figure S9a**) and performed *A. tumefaciens-*mediated transient expression assays in *N. benthamiana* leaves using the same conditions as Bartetzko and coworkers [69]. Under control conditions, SP_PR1_-GFP is secreted to compartments with acidic pH preventing it from fluorescing and we observed only a faint signal from the perinuclear endomembrane compartments (**Figure S9b**). GFP fluorescence signal intensity and distribution was altered by treatment with the secretion pathway inhibitor brefeldin A (BFA), and resulted in the formation of GFP-positive BFA bodies (**Figure S9b**). Co-expression of SPpr1-GFP with FLAG-REX2 did not affect GFP levels, while FLAG-REX3 enhanced GFP levels in perinuclear endomembrane compartments and resulted in a strong labelling of the cortical ER (**Figure 6c**). The ability of REX3 to retain GFP in endomembrane compartments suggest that this effector may promote infection by interfering with host secretion pathways.

### The TIPTOP promoter is activated at root tip infection sites

When screening our data for plant promoters responding early to *P. palmivora* attack we found Niben101Scf03747g00005, encoding a small secreted protein containing two repeats of a conserved SGPS-GxGH motif known from pathogen-associated molecular pattern (PAMP)-induced peptides (PIP/PIP-like; **Figure S11**) [32] to be one of the most strongly induced plant genes. To study the spatial distribution of its promoter activity we generated transgenic *N. benthamiana* plants expressing a promoter-*GFP:uidA* reporter fusion and challenged them with *P. palmivora* LILI-td [70] expressing a red fluorescent protein. Consistent with the transcriptomics data, histochemical GUS staining revealed a localised GUS signal at the tip of infected roots (**Figure 8**) only where zoospores had accumulated but not in uninfected roots. We therefore termed the gene *TIPTOP* (*Tip Induced Plant Transcript switched On by P. palmivora*). *TIPTOP* promoter activation is correlated with *P. palmivora* infection (**Figure 8c**). *P. palmivora*-triggered *TIPTOP* promoter activation was strongest adjacent to invasive hyphae as revealed by GFP confocal fluorescence microscopy (**Figure 8d**). In addition, the TIPTOP promoter was not activated by abiotic stresses (cold, heat and 1 M sodium chloride) and wounding, but weak activation was observed in root tips in response to flagellin (flg22) treatment (**Figure S14**). PlantPAN 2 [71] analysis of the *TIPTOP* promoter sequence identified various transcription factor binding motifs (**Table S5**). Taken together, these results suggest that *TIPTOP* is a root tip specific *P. palmivora*-induced promoter.

## Discussion

We utilised a dual transcriptomics approach coupled to a semi-automatic secretome annotation pipeline to study the interaction between *P. palmivora* and *N. benthamiana* roots. While the pathogen transcriptome undergoes remarkable shifts in expression patterns throughout the infection we see a steady response of the plant transcriptome with no detectable major shifts in sets of differentially expressed genes. We used our dataset to identify *P. palmivora* and *N. benthamiana* genes implicated in the interaction and characterised two conserved biotrophic *P. palmivora* effector proteins which confer enhanced infection susceptibility when expressed *in planta.* We show, that one of them, REX3, suppresses plant secretion processes. Surveying the set of early transcriptionally activated plant genes resulted in the identification of an *N. benthamiana* gene specifically induced at infected root tips and encoding a peptide with danger-associated molecular features.

### Dual transcriptomics and *de novo* assembly enables functional studies of unsequenced genomes

Dual transcriptomics captures simultaneous changes in host and pathogen transcriptomes [72,73] when physical separation of two interacting organisms is unfeasible. The diversity of plant pathogens often results in the absence of microbial reference genomes. This is particularly relevant for obligate biotrophic plant pathogens which cannot be cultivated separately from their host. Our established viable alternative, a *de novo* assembly of a plant pathogen transcriptome from separated mixed reads followed by an semi-automated annotation is thus applicable to a broader community. Taking advantage of the availability of the host reference genome, we separated *P. palmivora* reads from the mixed samples and combined them with reads from the *ex planta* samples to create a single *de novo* assembly for the pathogen transcriptome.

Assembly completeness in terms of gene content might be assessed based on evolutionary expectations, so that recovery of conserved genes serves as a proxy measure for the overall completeness (CEGMA [74] and BUSCO [55]). Our *P. palmivora de novo* assembly had sufficient read support (on average 76% reads mapping back), so we further probed it for the presence of BUSCOs. Since there is no specific oomycete set, we checked presence of 429 eukaryotic BUSCO genes and found 326 of them (76%). Lack of some BUSCO genes in our assembly might result from the fact that originally BUSCO sets were developed to estimate completeness of genomic assemblies and did not require expression evidence [55]. To verify this, we extended the same completeness analysis to existing *Phytophthora* genomes and transcriptomes and found that transcriptomes in general indeed contained fewer BUSCOs. Moreover, we found 35 eukaryotic BUSCO genes consistently missing from *Phytophthora* genomic assemblies. Therefore, a BUSCO-based completeness test for transcriptomes should be applied with caution within the *Phytophthora* genus, considering adjustments for expression support and uneven distribution of eukaryotic singlecopy orthologs. We propose that with an ever-growing body of oomycete genomic and transcriptomic data a specific set of benchmarking orthologs needs to be created to support *de novo* assemblies and facilitate studies of these economically relevant non-model plant pathogens [75].

So far, dual transcriptomics has only been used with limited time resolution and sequencing depth in plant-pathogenic oomycete studies [76,77]. Our study encompasses the full range of *P. palmivora* sequential lifestyle transitions occurring in *N. benthamiana* root, allowing reconstruction of comprehensive transcriptional landscape in both interacting organisms. We found three major waves of *P. palmivora* gene expression peaks that correlate with its major lifestyle transitions: 1) early infection and biotrophic growth inside host tissues; 2) switch to necrotrophy; 3) late necrotrophy and sporulation. Similar transcription dynamics following switches of life styles were previously described for the hemibiotrophic pathogens *Colletotrichum higginsianum* [78], *Phytophthora parasitica* [79] during *Arabidopsis* root infection and *P. sojae* upon infection of soybean leaves [80], though the exact timing of infection was different.

Interestingly, the *N. benthamiana* transcriptional response to infection does not mirror the observed significant shifts in infection stage specific *P. palmivora* gene expression. Instead it is characterized by steady induction and repression. High-resolution transcriptomics were applied to *A. thaliana* leaves challenged with *Botrytis cinerea* to untangle the successive steps of host response to infection [81]. However, in the absence of pathogen expression data, it is not possible to correlate these changes with changes in the pathogen transcriptome. It is likely that pathogen expression patterns are not useable to infer a link to corresponding plant responses.

The response of *N. benthamiana* roots to *P. palmivora* is characterised by an upregulation of genes associated with hormone physiology, notably ethylene through activation of ethylene response transcription factors (ERFs) and ACC synthase. Ethylene is involved in *N. benthamiana* resistance to *P. infestans* [82]. We also observed an induction of two PIN-like auxin efflux carriers. Suppression of the auxin response was associated with increased *A. thaliana* susceptibility to *P. cinnamomi* disease and was stimulated by phosphite-mediated resistance [83]. Interestingly, phosphite was also required for defense against *P. palmivora* [11]. We found upregulation of chitinases and endopeptidase inhibitors, such as Kunitz-type trypsin inhibitors, which are often induced by oomycete and fungal pathogens [84–86]. Induction of genes encoding O-glycosyl hydrolases is associated with cell wall remodelling while phenylalanine ammonia lyases (PAL) contribute to cell wall reinforcement by activation of lignin biosynthesis [87,88]. Upregulation of the trehalose biosynthesis pathway is associated with membrane stabilisation [89] and partially mitigates toxic effects of oxidative stress [90]. Upregulation of several enzymes of the mevalonate pathway suggest modulation of the biosynthesis of isoprenoids such as defense-associated phytoalexins as well as sterols. In particular, transcriptional repression of genes encoding sterol 4-alpha-methyl-oxidase 2-1 and C5 sterol desaturases suggest an attenuation of the brassinosteroid synthesis, while repression of genes with homology to sterol methyl transferase 2 point to a repression of the beta-sitosterol/stigmasterol branch. Conversely, induction of terpenoid synthases/epi aristolochene synthases points to a selective induction of the sesquiterpenes which contain defense associated phytoalexins such as capsidiol [91,92]. Finally, the *N. benthamiana* response to *P. palmivora* also includes upregulation of genes encoding late embryogenesis abundant (LEA) proteins as well as heat shock proteins. LEA proteins have been associated with the drought response [93,94] and upregulation of genes associated with water deprivation upon *Phytophthora* infection has been previously reported [77]. Conversely, downregulated genes were mostly associated with photosynthesis, cellulose biosynthesis and cell division. These results were consistent with previous reports [95,96].

### Analysing partial transcripts improved the predicted *P. palmivora* secretome

To study *P. palmivora* secreted proteins we developed a prediction and annotation pipeline tailored for signal peptide prediction based on ORFs derived from *de novo* assembled transcripts. Often a six-frame translation is utilised to identify candidate ORFs (Stothard, 2000; Lévesque et al 2010). However, we use a TransDecoder approach which enriches for the most likely translated ORF by integrating homology based matches to known Pfam domains and *Phytophthora* proteins. Compared to six-frame translation, this approach can result in partial ORFs which may lead to a mis-prediction of translation start sites and therefore signal peptides. So, we implemented a refinement step in our secretome prediction pipeline to rescue partial ORFs by finding the next likely translation start position and following the secretome prediction steps. This procedure allowed us to rescue an additional 611 ORFs including several which likely encode RXLR effectors, elicitins and cell wall degrading enzymes thus highlighting the importance of this additional step.

### Effector-guided resistance breeding potential

We identified two RXLR effectors that show high sequence conservation among *P. palmivora* isolates worldwide, suggesting they may represent core effectors that cannot be lost or mutated without a fitness cost for the pathogen [21]. As such, these effectors constitute valuable candidates to accelerate cloning of disease resistance (*R*) genes and effector-assisted deployment of resistance. This strategy has been used against *P. infestans* [22].

Our approach identified a potential AVR3a homolog in *P. palmivora* (PLTG_13552). The *P. infestans* AVR3a^KI^ allele confers avirulence to *P. infestans* isolates on *R3a*-expressing potatoes while the AVR3a^EM^ allele is not being recognised [63]. It will be interesting to study whether potato *R3a* or engineered *R3a* derivatives with a broader recognition spectrum [97,98] can be exploited to generate resistance towards *P. palmivora* in economically relevant transformable host plants. Additionally, *P. palmivora* proteins also harbour pep13-type MAMP motifs present in four transglutaminases and several nlp20-containing NLPs. While the pep13 plant receptor remains to be found, the receptor like kinase RLP23 has recently been identified as nlp20 receptor [99] with the potential to confer resistance even when transferred into other plant species. Introduction of RLP23 into *P. palmivora* host plants may thus be another strategy to engineer resistant crops.

### The *P. palmivora* effector REX3 inhibits plant secretion pathways

We found that REX3 interferes with host secretion, a common strategy of bacterial and oomycete pathogens [49,69]. Rerouting of the host late endocytic trafficking to the extrahaustorial membrane [41,100] and accumulation of the small GTPase RAB5 around haustoria [42] is well documented. Given that REX3 is almost invariant in *P. palmivora* it is likely that REX3 targets components of the secretory pathway which are conserved among diverse host species. Of the four functionally tested RXLR effectors the two most conserved ones (REX2, REX3) amongst *P. palmivora* isolates both conferred increased susceptibility. REX2 and REX3 therefore represent important targets for disease resistance breeding in tropical crops. It is possible that isolate-specific variants of REX1 and REX4 may provide a colonisation benefit only in hosts other than *N. benthamiana.*

### *P. palmivora* triggers expression of danger-associated molecular pattern peptides

Upon *P. palmivora* root infection 2’886 *N. benthamiana* genes were up and 3’704 genes down-regulated. Compared to previously studied root transcriptomes of responses to broad-host-range *Phytophthora* species [95,101] our data permitted the identification of early induced genes such as *TIPTOP,* a *P. palmivora*-responsive root tip promoter. An exciting future perspective is its exploitation for induced early resistance against *Phytophthora* root infections. This promoter also provides inroads to dissect early host cell responses to *P. palmivora,* when employed in combination with a cell sorting approach to generate samples enriched for infected cells.

The *TIPTOP* gene encodes a peptide with similarities to DAMP peptides [102]. The occurrence of two tandem repeats of a conserved sGPSPGxGH motif in the TIPTOP protein is reminiscent of the SGPS/GxGH motifs of PIP and PIPLs peptides [32,103] and the closest *Arabidopsis* homologs of TIPTOP, PIP2 and PIP3, are implied in responses to biotic stress.

Hou and coworkers showed that the PIP1 peptide is induced by pathogen elicitors and amplifies *A. thaliana* immune response by binding to the receptor-like kinase 7 (RLK7) [32]. Analysis of *in silico* data showed that PIP2 and PIP3 were activated upon *A. thaliana* infection by *Botrytis cinerea* or *P. infestans* [103]. By contrast to PIP1 and PIP2, the TIPTOP promoter is inactive under control conditions, suggesting it may undergo a different transcriptional regulation than the previously characterized *Arabidopsis* peptides.

## Conclusions

Dual transcriptomics represent a successful approach to identify transcriptionally regulated effectors as well as plant genes implicated in the root infection process. We found conserved MAMPs and effectors with similarity to known AVR proteins such as AVR3a which may harbour the potential for disease resistance engineering. We characterised two conserved RXLR effectors conferring enhanced susceptibility to root infection and confirmed interference with host secretion as a *P. palmivora* pathogenicity mechanism. Furthermore, the *P. palmivora* inducible TIPTOP promoter and the PIP2,3-like peptide are promising leads for engineering *P. palmivora* resistance. In summary, our findings provide a rich resource for researchers studying oomycete plant interactions.

## Methods

### Plant material and growth conditions

*N. benthamiana* seeds were surface sterilized for 3 min with 70 % ethanol and 0.05 % sodium dodecyl sulfate (SDS) and rinsed twice in sterile water. Seeds were cold-stratified for 2 days and sown on Murashige and Skoog (MS) medium (Sigma Chemical Company) supplemented with 20 g/L sucrose and 10 g/L agar. For *in vitro* susceptibility assays, two-week-old plants were transferred to square Petri dishes using the hydroponics system described elsewhere [104]. These dishes, each containing five plants, were then placed slanted for 2 weeks at 25°C under a 16-h photoperiod. For inoculations, zoospore suspension was added directly to the root compartment containing the liquid medium.

### *P. palmivora* growth conditions and *N. benthamiana* root inoculation

*P. palmivora* Butler isolate LILI (reference P16830) was initially isolated from oil palm in Colombia [70], and maintained in the *P. palmivora* collection at the Sainsbury Laboratory (Cambridge, UK). Transgenic *P. palmivora* LILI strain expressing KDEL-YFP [9] and tdTomato [70] have been previously described. *Phytophthora* growth conditions and the production of zoospores have been described elsewhere [10].

### Root inoculation and disease progression assays

For the investigation of effector dynamics during infection and activation of the TIPTOP promoter, we added 10^5^ *P. palmivora* zoospores to the liquid medium of Petri dishes containing 20-d-old plantlets grown as described already. Root infection assays were adapted from the *A. thaliana–P. parasitica* infection system described by Attard and coworkers [104]. One-week-old *N. benthamiana* seedlings were grown on hard (2 %) agar strips with roots immersed in 1/10th liquid MS medium for two weeks. Plates were then inoculated with 500 zoospores of *P. palmivora* LILI KDEL-YFP. Plants were scored on a daily basis using a disease index composed of five symptom extent stages (SES): healthy plants with no noticeable symptoms were given a SES value of 1. Plants with at least one wilted leaf were given a SES value of 2. Plants showing a brownish, shrunken hypocotyl were given a SES value of 3. Plants showing a brownish, shrunken hypocotyl and stem with multiple invaded or wilted leaves were given a SES value of 4. Finally, dead plants were given a SES value of 5. Statistical analyses of disease severity were based on Scheirer-Ray-Hare nonparametric two-way analysis of variance (ANOVA) for ranked data (*H*-test) [105].

### Quantitative reverse transcription-polymerase chain reaction (qRT-PCR) analyses

Total RNA was extracted from frozen, axenically grown mycelium with sporangia (sample MZ) and infected roots harvested at 3, 6, 18, 24, 30, 48 and 72 hours after inoculation (hai) using RNeasy Plant Mini Kit (Qiagen, USA). One microgram was reverse transcribed to generate first-strand cDNA, using the Roche Transcriptor First Strand cDNA Synthesis Kit according to the manufacturer’s instructions (Roche, Switzerland). Quality was assessed by electrophoresis on agarose gel. qRT-PCR experiments were performed with 2.5 μl of a 1:20 dilution of first-strand cDNA and LightCycler 480 SYBR Green I Master mix, according to the manufacturer’s instructions (Roche, Switzerland). Gene-specific oligonucleotides were designed with BatchPrimer3 software [106] (**Table S3**) and their specificity was validated by analyzing dissociation curves after each run. Genes encoding the *P. palmivora* orthologs of *P. parasitica* elicitor OPEL and a 40S ribosomal subunit S3A (WS21) were selected as constitutive internal controls for *P. palmivora* genes [107]. Genes encoding L23 (Niben101Scf01444g02009) and FBOX (Niben101Scf04495g02005) were selected as constitutive internal controls for *N. benthamiana* genes [108]. Three biological replicates of the entire experiment were performed. Gene expression was normalized with respect to constitutively expressed internal controls, quantified and plotted using R software.

### Plasmid construction

The vector pTrafficLights was derived from pK7WGF2 (Plant System Biology, Gent University, Belgium). A cassette containing the signal peptide sequence of *Nicotiana tabacum* pathogenesis-related protein 1 (PR-1; GenBank accession X06930.1) fused in frame with the green fluorescent protein (GFP) was obtained by PCR using primers SP-F/SP-R (**Table S3**) and ligated into pK7WGF2 using SpeI and EcoRI restriction enzymes. The AtUBQ10*pro*⸬DsRed cassette was amplified from pK7WGIGW2(II)-RedRoot (Wageningen University, Netherlands) using primers RedRoot-F/RedRoot-R (**Table S3**) and ligated into pK7WGF2 using XbaI and BamHI restriction enzymes.

The TIPTOP promoter (1230 bp, ending 46 bp before start codon) was PCR-amplified from *N. benthamiana* genomic DNA using primers TIPTOP-F2/TIPTOP-R2 (**Table S3**) and cloned into pENTR/D-Topo vector (Life Technologies Inc., Gaithersburg, Maryland, USA). The entry vector was then used for LR recombination (Life Technologies Inc., Gaithersburg, Maryland, USA) into expression vector pBGWFS7 (Plant System Biology, Gent University, Belgium).

### Transient *Agrobacterium tumefaciens* mediated expression

For transient expression of effectors in *N. benthamiana* leaves, *A. tumefaciens* cells (strain GV3101-pMP90) were grown overnight with appropriate antibiotics. The overnight culture was then resuspended in agroinfiltration medium composed of 10 mM MgCl_2_, 10 mM 2-(N-morpholino) ethanesulfonic acid (MES) pH 5.7 and 200 μM acetosyringone. Optical density at 600 nm (OD_600_) was then adjusted to 0.4 for transient expression of effectors. For secretion inhibition assays, effectors and pTrafficLights construct were mixed together in a 1:1 ratio to a final OD600 of 0.8. Agroinfiltrations were performed after 3-h-long incubation at 28°C using a syringe without a needle on the abaxial side of 5-week-old *N. benthamiana* leaves.

### Generation of transgenic *Nicotiana benthamiana*

*N. benthamiana* stable transformation was performed according to [109] with the following modifications: leaf discs were incubated in shoot-inducing medium (SIM) composed of 1X Murashige and Skoog (MS) medium supplemented with 2 % sucrose, 0.7 % agar, 50 mg/L kanamycin, 50 mg/L carbenicillin, 500 mg/L timentin and a 40:1 ratio of 6-benzylaminopurine (BAP) and 1-naphthaleneacetic acid (NAA). Emerging shoots were cut and transferred to root-inducing medium (RIM), which has same composition as SIM without BAP. After the first roots emerged, plantlets were transferred to soil and grown at 25°C under 16-h photoperiod.

### Histochemical staining for GUS activity

Transgenic *N. benthamiana* plantlets carrying pTIPTOP*pro*⸬GFP:GUS sequence were harvested 14 hours after inoculation and incubated in a staining solution containing 100 mM sodium phosphate pH 7.0, 0.1% (v/v) Triton X-100, 5 mM K_3_Fe(CN)_6_, 5 mM K_4_Fe(CN)_6_and 2 mM 5-bromo-4-chloro-3-indoxyl-β-D-glucuronid acid (X-gluc). Staining was carried out for 3 hours at 37°C. The plantlets were then washed with distilled water and observed with an AxioImager M1 epifluorescence microscope (Zeiss, Germany) equipped for Nomarski differential interference contrast (DIC).

### Confocal microscopy

Confocal laser scanning microscopy images were obtained with a Leica SP8 laser-scanning confocal microscope equipped with a 63x 1.2 numerical aperture (NA) objective (Leica, Germany). A white-light laser was used for excitation at 488 nm for GFP visualization, at 514 nm for YFP visualization and at 543 nm for the visualization of tdTomato. Pictures were analysed with ImageJ software (http://imagej.nih.gov/ij/) and plugin BioFormats.

### Library preparation and sequencing

*N. benthamiana* and *P. palmivora* mRNAs were purified using Poly(A) selection from total RNA sample, and then fragmented. cDNA library preparation was performed with the TruSeq^®^ RNA Sample Preparation Kit (Illumina, US) according to the manufacturer’s protocol. cDNA sequencing of the 13 samples (MZ, infected *N. benthamiana* root samples and uninfected *N. benthamiana* plants) was performed in 4 lanes of Illumina NextSeq 2500 a in 100 paired end mode. Samples were de-multiplexed and analyzed further. mRNAs from additional samples of a short leaf time course (*P. palmivora* mycelium, *N. benthamiana* leaves 2 dai and *N. benthamiana* leaves 3 dai) were purified using Poly(A) selection from total RNA sample. cDNA libraries were prepared using NEBNext^^®^^ RNA library preparation kit (New England Biolabs, UK) according to the manufacturer’s protocol and sequenced on Illumina GAII Genome Analyzer in a 76 paired end mode in 3 separate lanes. Reads obtained from these three samples were used for *P. palmivora de novo* transcriptome assembly only. The raw fastq data are accessible at http://www.ncbi.nlm.nih.gov/sra/ with accession number SRP096022.

### *De novo* transcriptome assembly

In order to capture the full complexity of the *P. palmivora* transcriptome we pooled all the samples potentially containing reads from *P. palmivora* (**Figure 2**): eight mixed (plant-pathogen, combining leaf and root infections), one exclusively mycelium and one mixed mycelium-zoospores sample. Initial read quality assessment was done with FastQC (Babraham Bioinformatics, Cambridge, UK). Adaptors were removed using CutAdapt [110]. To exclude plant reads from the library, raw paired-reads were first aligned to *N. benthamiana* reference genome (v1.01) using Tophat2 [111]. Unmapped reads (with both mates unmapped) were collected with samtools (samtools view -b -f 12 -F 256), converted to fastq with bedtools and processed further. To estimate the level of residual contamination by plant and potentially bacterial reads, the resulting set of reads was subjected to FastQ Screen against the UniVec database, all bacterial and archaeal sequences obtained from RefSeq database, all viral sequences obtained from RefSeq database, *N. benthamiana* genome (v1.01), and subset 16 oomycete species (mostly *Phytophthora* species). Since the above test revealed substantial residual contamination by *N. benthamiana* reads, an additional round of bowtie2 alignment directly to *N. benthamiana* transcriptome [53] was performed followed by FastQ Screen. Reads, not aligned to *N. benthamiana* genome and transcriptome were further subjected to quality control using Trimmomatic (minimum read length = 60). The quality parameters for the library were assessed using FastQC. The total of ∼190 M filtered reads were subjected to *de novo* assembly with Trinity (trinity v2.1.1) on a high-RAM server with minimal k-mer coverage = 2 and k-mer length = 25. *In silico* read normalization was used due to the large number of input reads, in order to improve assembly efficiency and to reduce run times [57]. The resulting assembly was additionally checked for plant contamination using blastn search against plant division of NCBI RefSeq genomic database. Trinity genes having significant sequence similarity (e-value threshold ≤10-5) to plant sequences were removed from the resulting transcriptome. The final version of assembly included trinity genes with sufficient read support.

### *De novo* assembly statistics and integrity assessment

General statistics of the assembly were determined using the ‘TrinityStats.pl’ script provided with Trinity release and independently using Transrate (http://hibberdlab.com/transrate/) and Detonate (http://deweylab.biostat.wisc.edu/detonate/) tools. Assembly completeness was estimated using the eukaryotic set of BUSCO profiles (v1) [55]. BUSCO analysis was performed for the full transcriptome assembly and for the reduced assembly, obtained after retaining only the longest isoform per trinity gene. BUSCO genes missing from the assembly were annotated with InterProsScan based on the amino acid sequences emitted from the corresponding hmm profile (‘hmmemit’ function from hmmer package, http://hmmer.org/). Overall expression support per assembled transcript was performed after transcript abundance estimation. Trinity genes with TPM≥ 1 in at least 3 samples were considered further.

### Protein prediction and annotation

ORFs were predicted using TransDecoder software [57]. At the first step ORFs longer than 100 aa were extracted. The top 500 longest ORFs were used for training a Markov model for coding sequences, candidate coding regions were identified based on log-likelihood score. Additionally all the ORFs having homology to protein domains from the Pfam database and/or *P. sojae*, *P. parasitica*, *P. infestans* and *P. ramorum* protein sequences downloaded from Uniprot database (accession numbers: UP000005238, UP000006643, UP000002640, UP000018817) were also retained (blastp parameters: max_target_seqs 1 -evalue 1e-5).

### Secretome prediction

For the automatic secretome prediction a custom script was written, employing steps taken for *P. infestans* secretome identification [16]. Predicted proteins were subsequently submitted to SignalP 2.0 (Prediction = ‘Signal peptide’), SignalP 3.0 (Prediction = ‘Signal peptide’, Y max score ≥ 0.5, D score ≥ 0.5, S probability ≥ 0.9), TargetP (Location = ‘Secreted’) [112] and TMHMM (ORFs with transmembrane domains after predicted signal peptide cleavage site were removed) [113]. Finally, all proteins with terminal ‘KDEL’ or ‘HDEL’ motifs were also removed, as these motifs are known to be ER-retention signals [114]. Exact duplicated sequences and substrings of longer ORFs were removed to construct non-redundant set of putative secreted proteins. Taking into account possible fragmentation of *de novo* assembled transcripts a custom python script (M-slicer) was developed to rescue partial proteins with mis-predicted CDS coordinates. It takes as an input all the partial translated ORFs, which were not predicted to be secreted initially and creates a sliced sequence by finding the position of the next methionine. The M-sliced proteins were subjected to the same filtering step as was done with the initial secretome. The same script, omitting the M-slicer refinment, was used to systematically predict *N. benthamiana* genes encoding putative secreted proteins.

### Secretome annotation

To annotate putative secreted proteins a complex approach was used, combing several lines of evidence: 1) blastp search against GenBank NR database with e-value ≤10-6; 2) InterProScan (version 5.16) search against databases of functional domains (PANTHER, Pfam, Coils, Gene3D, SUPERFAMILY, SMART, PIRSF, PRINTS) with default parameters [115]; 3) RXLR and EER motif prediction using regular expressions; 4) WY motif prediction based on WY-fold HMM by hmmsearch function from HMM3 package (http://hmmer.org/); 5) LxLFLAK and HVLVVVP motif predictions based on HMM model build on sequences of known CRN effectors; 6) NLS motif prediction by NLStradamus [116] (version 1.8, posterior threshold = 0.6) and PredictNLS [117] with default parameters. The TribeMCL algorithm was used to cluster predicted putative secreted proteins with signal peptide and after signal peptide cleavage (mature proteins). The tribing results were used as a soft guidance for functional annotation (proteins belonging to the same tribe are likely to have the same function). All obtained data were aggregated in the **Supplementary Dataset: Annotated *P. palmivora* secretome**. Functional categories were assigned based on manual curation of the resulting table. ‘Hypothetical’ category was assigned to proteins either having similarity to only hypothetical proteins or when the top 20 hits of blastp output did not show consistency in terms of distinct functional categories. Proteins having significant sequence similarity to ribosomal, transmembrane proteins or proteins with known intracellular localization (e.g. heat shock proteins) and/or having respective domains identified by InterProScan were marked as false predictions. A contamination category was assigned for proteins with significant sequence similarity (revealed by blastp) to amino acid sequences from phylogenetically distant taxa (e.g. plants or bacteria). Entries marked as both ‘false prediction’ or ‘contamination’ were excluded from the final secretome.

### Transcriptome annotation

All the remaining predicted proteins were annotated by scanning against InterProScan databases of functional domains (version 5.16-55) and by performing blastp search against GenBank NR database (download date: 06.01.2016) and published reference *Phytophthora* genomes.For transcripts without predicted ORFs blastn search against the GenBank NR database was performed, and the top hit with e-value ≤ 10-5 was reported (**Supplementary Dataset: Whole**_**transcriptome**_**expression**_**TMM**_**TPM**_**normalised**_**filtered**_**PLTG**).

### Expression analysis

Initial reads after quality control were separately aligned aligned back to the *P. palmivora de novo* transcriptome assembly and *N. benthamiana* reference transcriptome. Alignment-based transcript quantification was done using RSEM (version: RSEM-1.2.25, (http://deweylab.github.io/RSEM/)[118]. For *P. palmivora* quantification was performed on ‘trinity gene’ level. For within-sample normalisation TPMs were calculated. Between-sample normalisation was done using trimmed means approach (TMM) [119]. TMM-normalised TPMs were reported for both *P. palmivora* and *N. benthamiana.* PCA-analysis was performed on the log-transformed TPM values and visualized in R with the help of “ggplot2” [120] and “pheatmap” [121] packages. Overlap between groups of genes identified in the PCA analysis was visualised with “Vennerable” package [122]. Differentially expressed genes were identified with edgeR Bioconductor package [119] following pair-wise comparisons between all the samples. The dispersion parameter was estimated from the data with the estimateDisp function on reduced datasets: for *P. palmivora* we combined close time points (based on PCA analysis) and treated them as pseudo-replicates; for *N benthamiana* common dispersion was estimated based on 6 uninfected plant samples, treating them as replicates. The resulting common dispersion values of 0.15 and 0.1 were used for *P. palmivora* and *N. benthamiana* analysis, respectively. Most differentially expressed genes (log2(fold change) ≥2 and p-value ≤103 were used to perform hierarchical clustering of samples. Heatmaps for the most differentially expressed genes were generated using R “cluster” [123], “Biobase” [124] and “qvalue” packages. For the final heatmaps TPMs were log2-transformed and then median-centered by transcript. Plant samples were centered according to the full set of mock and infected sample. Temporal clustering of expression profiles was done with fuzzy clustering (Mfuzz Bioconductor package) [68] to adopt gradual temporal changes of gene expression in the course of infection. GO-enrichment analysis was done with the help of “topGO” Bioconductor package [125]. Gene universe was defined based on *N. benthamiana* genes having expression evidence in our dataset (having TPM ≥ 1 in at least 3 samples). For the enrichment analysis exact Fisher test was used and GO-terms with p-values ≤ 05 were reported. ReviGO [126] was used to summarize the resulting significant GO-terms and reduce redundancy.

## Abbreviations

DEG: differentially expressed genes
CRN: crinkler effector
LFC: log fold change
FDR: false discovery rate
ORF: open reading frame
TPM: transcripts per million
TMM: trimmed mean of m-values
PCA: principal component analysis
BUSCO: benchmarking universal single-copy orthologs
REX: putative RXLR-effector expressed
NPP: necrosis-inducing *Phytophthora* protein
SCR: small cysteine-rich peptides
PLTG: *Phytophthora palmivora* transcribed gene
PI: protease inhibitor
TM: transmembrane domain
SP: signal peptide
SES: symptoms extend stage
aa: amino acid
hai: hours after inoculation
dai: days after inoculation

## Declarations

### Ethics approval and consent to participate

Not applicable

### Consent for publication

Not applicable

### Availability of data and materials

The raw fastq files are available in the SRA archive under SRP096022 accession number.

### Competing interests

The authors declare that they have no competing interests

### Funding

This work was supported by the Gatsby Charitable Foundation (RG62472), by the Royal Society (RG69135), and by the European Research Council (ERC-2014-STG, H2020, 637537).

### Authors’ contributions

KF, CQ and MD generated constructs. TY and EE generated *N. benthamiana* transgenics. FT and EE characterized transgenics. MD, EE and TH obtained microscopic data. AG performed bioinformatics analysis. EE, AG and SS analyzed the data and wrote the manuscript.

## Acknowledgements

We acknowledge the experimental and annotation help by Abhishek Chatterjee and Schornack lab. We would like to thank Mike Coffey and Joe Win for provision of pathogen isolates and list of oomycete RXLRs. We are indebted to Diane Saunders, Liliana Cano, Jodie Pike and Sophien Kamoun for generating leaf transcriptome sequences. We would like to thank Ruth Le Fevre and Stuart Fawke for proof-reading the manuscript.

## Supporting information

**Figure S1** – BUSCO genes missing from available *Phytophthora* genomes and transcriptomes.

**Figure S2** – Amino acid sequence alignment of PLTG_13552 and *P. infestans* AVR3a^EM^.

**Figure S3** – Number of DEGs between infection time points.

**Figure S4** – Validation of dynamic behavior of *P. palmivora* DEGs by qRT-PCR.

**Figure S5** – PCR detection of REX1-4 effectors in *P. palmivora* isolates.

**Figure S6** – Amino acid sequence logos for REX1-4 effectors.

**Figure S7** – Habitus of *N. benthamiana* transgenics used in this study.

**Figure S8** – Subcellular localisation of GFP:REX1-4 proteins in *N. benthamiana* leaves.

**Figure S9** – Structure of pTrafficLights construct and secretion inhibition assays.

**Figure S10** – Validation of *N. benthamiana* DEGs by qRT-PCR.

**Figure S11** – Amino acid sequence alignment of TIPTOP and similar *N. benthamiana* sequences with *A. thaliana* prePIPL1, prePIP1 and prePIP2.

**Figure S12** – Induction of TIPTOP promoter in response to biotic and abiotic stresses.

**Table S1** – Sequencing and mapping statistics in RNA-seq samples containing *P. palmivora*

**Table S2** – BUSCO genes missing from genomes and transcriptomes of *Phytophthora* genus. BUSCO genes were Annotated using InterProScan based on sequences emitted from HMM profiles.

**Table S3** – Primers used in this study

**Table S4** – *P. palmivora* isolates

**Table S5** – PlantPAN analysis of TIPTOP promoter sequence

**Supplementary Dataset 1** - ***P. palmivora de novo* transcriptome assembly.** Assembly was performed using trinity 2.1.1 software. CDS and corresponding mRNA and amino acid sequences were predicted using Transdecoder with additional homology-based filters.

Tar archive contains 4 files:

LILI_transcriptome_v5_converted.fasta - final version of P. palmivora transcriptome

LILI_transcriptome_v5.transdecoder.cds.fasta - Transdecoder-predicted CDS

LILI_transcriptome_v5.transdecoder.mRNA.fasta - mRNAs for predicted CDS

LILI_transcriptome_v5.transdecoder.pep.fasta - amino acid sequences

**Supplementary Dataset 2** - ***N. benthamiana* expression table. Raw counts were normalised within and between samples**. TMM-normalised TPMs were reported. Functional annotation provided with 1.01 version of N. benthamiana genome was used: (ftp://ftp.solgenomics.net/genomes/Nicotiana_benthamiana/annotation/Niben101/).

**Supplementary Dataset 3** - ***P. palmivora* expression table.** Raw counts were normalised within and between samples. TMM-normalised TPMs were reported. Not manually curated high throughput annotation (as described in Material and Methods) is provided.

**Supplementary Dataset 4** - **GO**-**enrichment for *N. benthamiana* genes up and downregulated during *P. palmivora* infection.** GO-enrichment was done with TopGO Bioconductor package. Classic Fisher test was used, only GO-terms with p-value < 0.05 reported.

**Supplementary Dataset 5 - Manually curated *P. palmivora* secretome.**

